# BaGPipe: an automated, reproducible, and flexible pipeline for bacterial genome-wide association studies

**DOI:** 10.1101/2025.02.28.640835

**Authors:** Kuangyi Charles Wei, Beth Blane, Jacqueline Toussaint, Sandra Reuter, Michelle S. Toleman, Estee Torok, Sharon J. Peacock, Ewan M Harrison, Dinesh Aggarwal, William Roberts-Sengier

## Abstract

Microbial genome-wide association study (GWAS) tools often require manual data processing steps, lack comprehensive workflows, and are limited by scalability issues, thus hindering the exploration of bacterial genetic traits. To address these challenges, we developed BaGPipe, an automated and flexible bacterial GWAS pipeline built using Nextflow and incorporating Pyseer for association analysis. BaGPipe integrates all essential components of a bacterial GWAS—spanning pre-processing, statistical analysis, and downstream visualisation—into a unified workflow that is reproducible and easy to deploy across diverse computational environments. BaGPipe was validated on a publicly available dataset of *Streptococcus pneumoniae* whole-genome sequences, and reproduced published findings with improved computational efficiency. BaGPipe was then applied to a dataset of *Staphylococcus aureus* whole-genome sequences, successfully identifying known and novel antibiotic resistance associations. By offering an accessible, efficient, and reproducible platform, BaGPipe accelerates bacterial GWAS and facilitates deeper exploration into the genetic underpinnings of phenotypic traits.

**Impact Statement:** The increasing availability of bacterial genome sequences has created an opportunity for robust, reproducible tools to facilitate the discovery of novel genotype-phenotype associations. Despite the demonstrated utility of genome-wide association studies (GWAS) in identifying genetic determinants of disease, toxicity and antibiotic resistance, existing tools for bacterial GWAS often involve fragmented workflows requiring extensive manual intervention, limiting their adoption and reproducibility. Here, we introduce BaGPipe, a fully integrated bacterial GWAS pipeline that automates pre-processing, statistical analysis, and visualisation, thereby streamlining the entire workflow. With its flexibility, scalability, and ease of use, BaGPipe makes bacterial GWAS more accessible to researchers, enabling faster and more reliable insights into microbial genetics. This is an important step towards overcoming the computational and logistical barriers that have constrained bacterial GWAS, ultimately accelerating research into microbial evolution, resistance mechanisms, and the genetic basis of other key phenotypic traits.

**Data Summary:** BaGPipe is freely available at https://github.com/sanger-pathogens/BaGPipe. The *Streptococcus pneumoniae* input dataset is available from the Pyseer tutorial (https://pyseer.readthedocs.io/en/master/tutorial.html#). The *Staphylococcus aureus* sequencing assemblies can be sourced from their ERS accession numbers provided in supplementary data. The reference assemblies, listed in the supplementary, can be sourced from NCBI.

## Introduction

Genome-wide association studies (GWAS) have become an essential tool for uncovering the genetic basis of phenotypic traits across diverse organisms, particularly for investigating mechanisms of pathogenicity and resistance [1–5]. However, bacterial GWAS faces unique challenges due to horizontal gene transfer, complex population structures, and extensive genomic diversity, which complicate accurate genotype-phenotype associations [6–9]. Current bacterial GWAS tools often exacerbate these difficulties by requiring fragmented workflows, extensive manual intervention, and lacking scalability and reproducibility.

Executing bacterial GWAS requires analysing large genomic datasets to identify associations with phenotypes such as antimicrobial resistance (AMR), while accounting for factors like population structure and statistical power [10]. Current bacterial GWAS tools fall into three categories [9, 11, 12]: phylogenetic, non-phylogenetic, and machine learning approaches (See **Supplementary Table 1**). Phylogenetic tools like Scoary [13] and TreeWAS [14] rely on well-defined phylogenetic structures and are suitable for datasets where recombination can be mitigated but are less practical for highly diverse species or large datasets. Non-phylogenetic tools such as Bugwas [15] and SEER [16] adapt concepts from human GWAS to develop phylogeny-independent methods. While computationally efficient and scalable, they face challenges like elevated false-positive rates due to insufficient correction for population structure.

Among non-phylogenetic tools, Pyseer [17] effectively addresses population structure and mitigates false positives by incorporating a linear mixed model [18]. By leveraging dimensionality reduction techniques like multidimensional scaling (MDS) and employing k-mer based association studies, Pyseer has emerged as a standard tool in the field due to its robustness and speed, especially in identifying genetic variants with small effect sizes [12, 19–22].

Despite these advancements, executing bacterial GWAS analyses remains hindered by significant technical challenges. Pyseer and similar tools [23–25] lack integrated solutions for crucial pre-processing steps—such as genome annotation, phylogenetic inference, or distance matrix generation—and downstream analyses like result visualisation and annotation (**Figure 1**). These steps demand expertise across multiple bioinformatics tools, each with its own dependencies and idiosyncrasies, leading to an increased risk of error, wasted computational resources, and ultimately, a diversion from biological interpretation towards technical troubleshooting. Existing pipelines, such as bacterialGWAS [26], DBGWAS [23] and GEMMA kmer_pipeline [27], attempt to address specific aspects of bacterial GWAS but remain limited by their narrow scope, fragmented workflows, or lack of scalability. A recent Snakemake pipeline, microGWAS [28], offers a more comprehensive workflow but it is unclear how flexible and scalable it is for diverse bacterial datasets.

**Figure 1:**
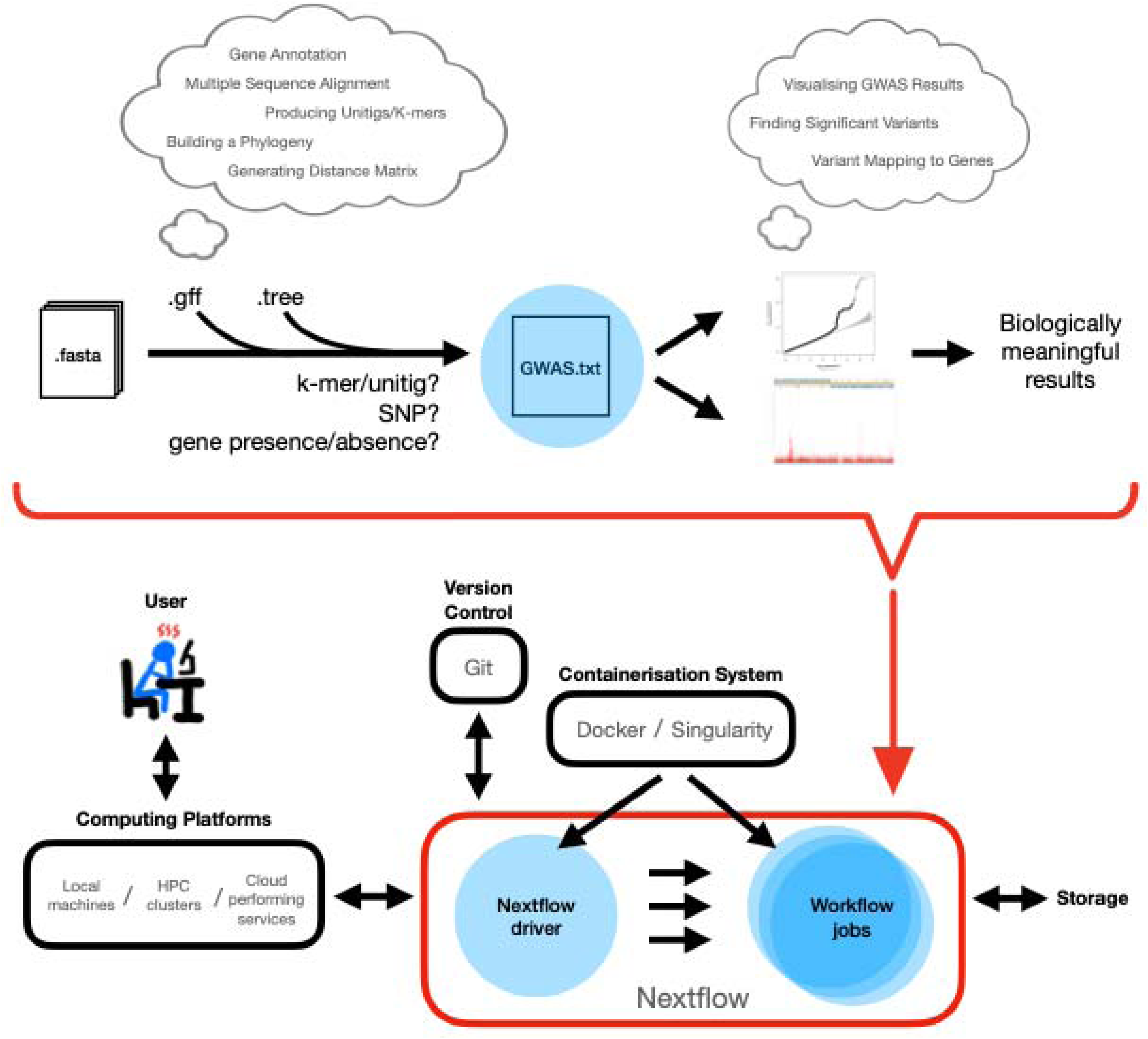
Schematic of the problem addressed and the solution by BaGPipe. Top: A comprehensive bacterial GWAS workflow involves miscellaneous pre-processing steps and various post-processing analyses, yet most bacterial GWAS tools only achieve the association study (shown by a blue circle). Bottom: The objective is to compile all relevant tools into a Nextflow pipeline, which can be easily deployed by users across computing platforms. The quantile-quantile plot (Q-Q plot) and the Manhattan plot are taken from the Pyseer tutorial [29].

To address these unmet needs, we present BaGPipe, a highly scalable and reproducible pipeline built on the workflow manager Nextflow [30] (**Figure 1**). BaGPipe fully leverages Pyseer for bacterial GWAS analysis while encompassing the entire GWAS workflow. We validate BaGPipe by replicating previously published findings in a well-characterised *Streptococcus pneumoniae* dataset [29, 31, 32] and further examine its utility by exploring novel genetic associations in a clinically relevant *Staphylococcus aureus* genomic dataset [33].

## Methods

### Pipeline Overview

BaGPipe is a bacterial GWAS pipeline designed to integrate a wide array of bioinformatics tools into a unified workflow that can efficiently handle pre-processing steps to post-processing analysis and visualisation. The pipeline was developed to address computational challenges and facilitate genome-wide association studies on bacterial datasets in a reproducible and user-friendly manner. The pipeline utilises Nextflow [30] for workflow management, ensuring compatibility across diverse computational environments while optimising resource usage. A step-by-step tutorial and comprehensive documentation are available online (https://github.com/sanger-pathogens/BaGPipe).

BaGPipe supports different entry points to provide maximum flexibility for users (**Figure 2**), allowing for the integration of their own input data, such as phylogenetic trees or variant call formants (VCFs). This feature allows BaGPipe to accommodate diverse user requirements, reducing the burden of having to repeat computationally intensive pre-processing tasks.

**Figure 2:**
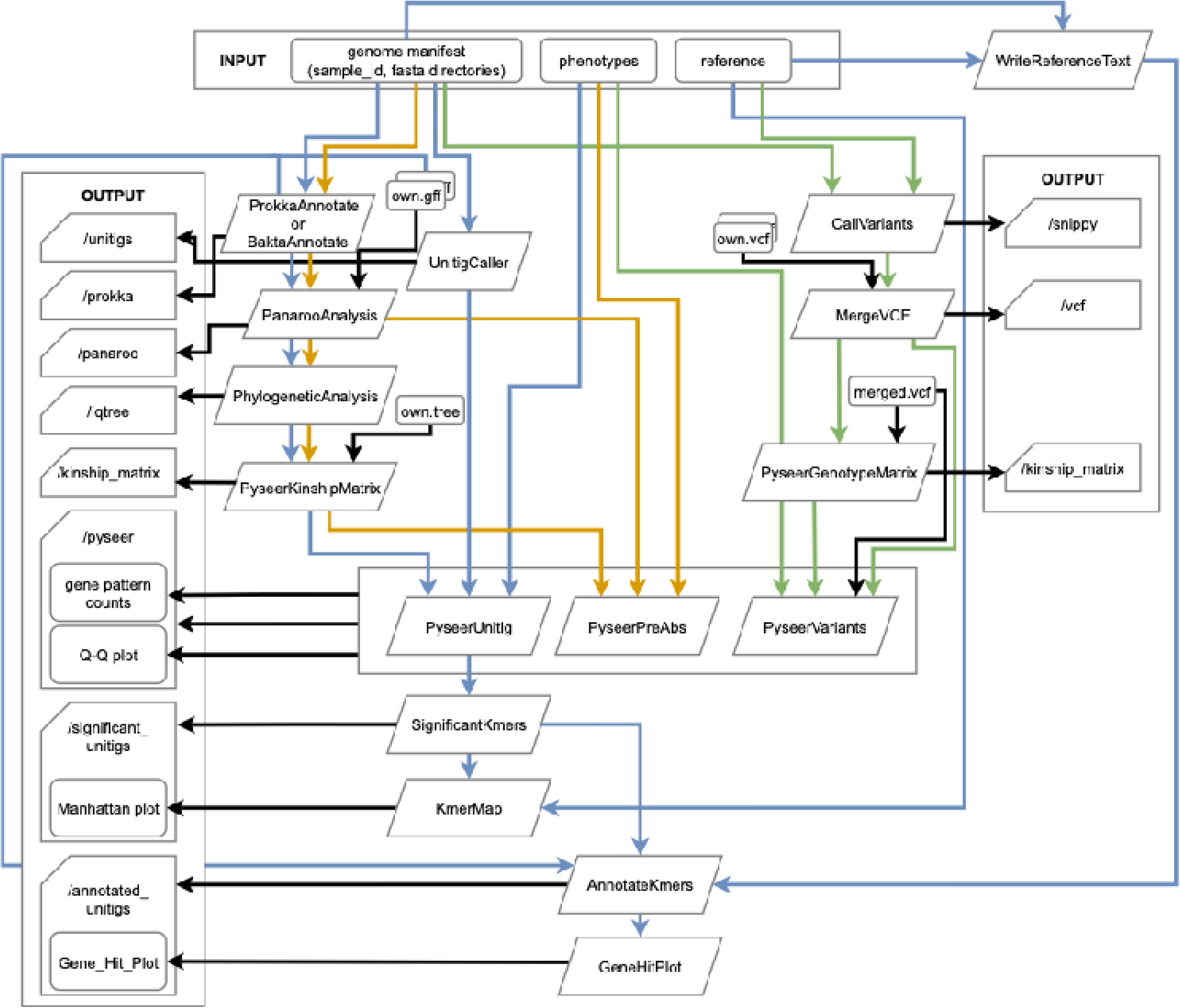
Overview of the BaGPipe pipeline. Input files are displayed on the top and output files on the sides. Alternative input files (GFF file, phylogenetic tree file, or VCF file) can be provided using appropriate option parameters. In a comprehensive analysis using the unitig mode, there will be eight output folders. The colour defines the genotype mode of analysis (blue: unitig/k-mer; yellow: gene presence and absence; green: short variations, including SNPs and Indels).

### Pipeline Implementation and Reproducibility

To ensure ease of use and reproducibility, BaGPipe was developed using Docker and Nextflow [30]. Docker standardises the software environment, encapsulating all essential tools and specific versions into a single container for consistency across platforms. Nextflow manages the modular pipeline, allowing users to execute the workflow with a single command while specifying input files and analysis settings:

\# install pipeline
nextflow pull sanger-pathogens/bagpipe
\# run pipeline
nextflow run sanger-pathogens/bagpipe –help

For high-performance computing (HPC) environments, BaGPipe supports the Load Sharing Facility (LSF) executor, enabling efficient parallel processing by dispatching each task as an independent job. It dynamically allocates memory and CPU based on task needs and can escalate resource requests if a job fails due to underestimation.

### User Accessibility and Configuration

The pipeline includes a default configuration file with all the key options and parameters necessary for running BaGPipe. This configuration is easily modifiable: non-experienced users need only adjust key parameters, while advanced users can modify any parameter by editing the source code or profile configuration. For example, the configuration can be easily extended to other Distributed Resource Managers (DRMs), such as Slurm. This ensures that BaGPipe remains both accessible and highly adaptable to meet the needs of varied research projects.

### Pre-processing

In its most comprehensive mode, BaGPipe starts with genome assemblies and automates pre-processing, including the generation of k-mers or unitigs, genome annotation, pangenome analysis, and phylogenetic analysis. This pre-processing stage is supported by Prokka (v.1.14.6) [34] and Bakta (v.1.10.2) [35] for annotation, Panaroo (v.1.5.1) [36] for constructing pangenomes and creating a multiple sequence alignment of core genes, and IQ-TREE (v.2.2.6) [37] for generating a core genome phylogeny. The output from these steps provides the essential elements for subsequent analysis, such as annotated assemblies, a core phylogenetic tree, and a distance matrix that reflects the genetic relationships within the dataset.

### Association Analysis

BaGPipe employs Pyseer (v.1.3.11) [17], a Python implementation of SEER [16], to conduct the association study, leveraging a linear mixed model to identify genotype-phenotype relationships. The typical bacterial GWAS requires three core types of inputs: genotypes, phenotypes, and an interaction or population structure correction. In BaGPipe, users can conduct association analyses using different genotyping approaches, including unitigs, k-mers, SNPs and indels, which provide flexibility in how genomic variation is represented.

The linear mixed model in Pyseer controls for population structure using kinship matrices derived from phylogenetic analysis. Alternatively, a pairwise distance matrix can be generated from genome assemblies using Mash [38], which is then employed to manage population stratification. BaGPipe supports Pyseer’s k-mer/unitig-based association study, which is recognised as a best practice for bacterial GWAS due to its capacity to handle high genetic diversity while controlling for population structure [29]. This k-mer/unitig approach aids in controlling population structure by providing a comprehensive genetic representation of each isolate, capturing all types of genetic variations—including SNPs, indels, and structural variants. By using k-mers/unitigs to quantify genetic similarities among isolates, a more accurate kinship matrix can be constructed, reflecting the true genetic relationships based on the presence or absence of k-mers across genomes.

Q-Q plots are generated to visualise the *p*-value distribution against the null hypothesis, ensuring that significant associations are not artefacts of confounding factors, such as population structure, phylogenetic relatedness, horizontal gene transfer, or technical biases. Additionally, as a good practice inherited from using Pyseer, BaGPipe provides a count of genotype patterns to guide users on whether corrections for multiple testing are necessary, if k-mers or unitigs are used as the genotyping approach.

### Post-processing and Visualisation

BaGPipe facilitates intuitive analysis by automatically processing significant unitigs or other genomic elements identified during the association analysis. It generates Manhattan plots to visualise the genomic locations of significant associations and annotates these markers using both reference and draft assemblies. The output includes a "gene_hits.tsv" file summarising significant unitigs and their gene annotations, providing insights into potential biological functions.

BaGPipe enriches the final output by annotating genes in the neighbourhood of significant markers, which aids in biological interpretation. Additionally, a default R script (adapted from the documentation of Pyseer [29]) is provided to allow users to create custom visualisations of the data, further simplifying downstream exploration and interpretation.

### Implementation of BaGPipe on the *Streptococcus pneumoniae* dataset

To assess BaGPipe’s ability to identify resistance-associated variants, we applied it to a well-characterised *S. pneumoniae* dataset (*n* = 616) on beta-lactam resistance. This dataset was chosen as a benchmark due to its prior use in bacterial GWAS [29, 31, 32], allowing for comparisons with existing methods like Pyseer. Different to the Pyseer tutorial [29], BaGPipe was run with unitigs instead of k-mers to reduce sequence redundancy while maintaining the integrity of the analysis. For inputting to BaGPipe, we made the required genome manifest CSV file indicating the paths to all assemblies. To conduct significant unitig analysis, we also made a reference manifest CSV file. BaGPipe was executed with the LSF executor.

### Implementation of BaGPipe on the *Staphylococcus aureus* dataset

To evaluate BaGPipe’s performance in a different genomic context, we applied it to a methicillin-resistant *S. aureus* (MRSA) dataset [33], aiming to identify genetic determinants of antibiotic resistance in a real-world genomic surveillance setting. To quality-control the assemblies, we ran the FASTA files (*n* = 520) through bacQC (v.1.2, https://github.com/avantonder/bacQC), which trimmed sequences using fastp [39]. Inspection on the output QC file confirms a slight improvement in the quality of the data, with fewer adapter sequences and PCR duplicate artifacts. From the Kraken2 [40] and Bracken [41] reports, two obviously contaminated reads were identified and excluded from the data for further analysis (see supplementary). Next, we forwarded the FASTQ files (*n* = 518) to assembleBAC (v.1.2, https://github.com/avantonder/assembleBAC). It was observed that the smallest N50 is 72.2 Kbp and all 518 assemblies have length 2.8-3.0 Mbp. We discarded samples with less than 90% reads matching to *S. aureus* and those with <30x coverage from onward analyses, as well as removing assemblies with an N50 value <10 Kbp, length of less than 2.6 Mbp or greater than 3.0 Mbp, or with a spuriously high number of contigs. No other samples were excluded based on these criteria.

We prepared the appropriate manifest files and other input files and executed BaGPipe using the LSF executor. Analyses were conducted to identify the genetic basis of resistance to eight antibiotics: oxacillin, ciprofloxacin, erythromycin, fusidic acid, clindamycin, tetracycline, gentamicin, and mupirocin. Antibiotic resistance in the *S. aureus* dataset was recorded as binary. Details of isolate collection, whole-genome sequencing and antimicrobial susceptibility testing are available in the published study [33].

To compare our results, we ran AMRFinderPlus (v.3.11.18) [42] for prediction of antimicrobial resistance genes from known databases. For each of the gene hits from BaGPipe, we found matches in the AMRfinder results in all genomes and counted the number of matches.

## Results

### Validation of BaGPipe by Reproducing Analyses from Pyseer

We validated BaGPipe on 616 *S. pneumoniae* isolates, the dataset used in the Pyseer tutorial [29, 31, 32]. This dataset has previously been used to identify genes associated with penicillin resistance, providing a benchmark for bacterial GWAS. By analysing the identical isolates, we aimed to assess BaGPipe’s ability to replicate these findings and evaluate its performance against established methods.

BaGPipe successfully identified 337,885 unique unitig patterns from a total of 740,945 unitigs tested. Recognising that many unitigs are highly correlated due to sequence overlap, we determined a significance threshold of 1.48E-07 using a Bonferroni correction based on the number of unique unitig patterns (0.05/337,885). This approach aligns with the Pyseer tutorial, which suggests adjusting the significance threshold based on the effective number of independent tests. We then applied this threshold to filter significant unitigs—those with *p*-values below 1.48E-07 were considered significant. By using this adjusted threshold, we effectively controlled for multiple testing without being overly conservative, ensuring that the identified associations are statistically robust. The Q-Q plot generated by BaGPipe closely resembled that produced in the Pyseer tutorial (**Figure 3**), demonstrating consistency in managing population structure and ensuring the absence of poorly controlled confounders.

**Figure 3:**
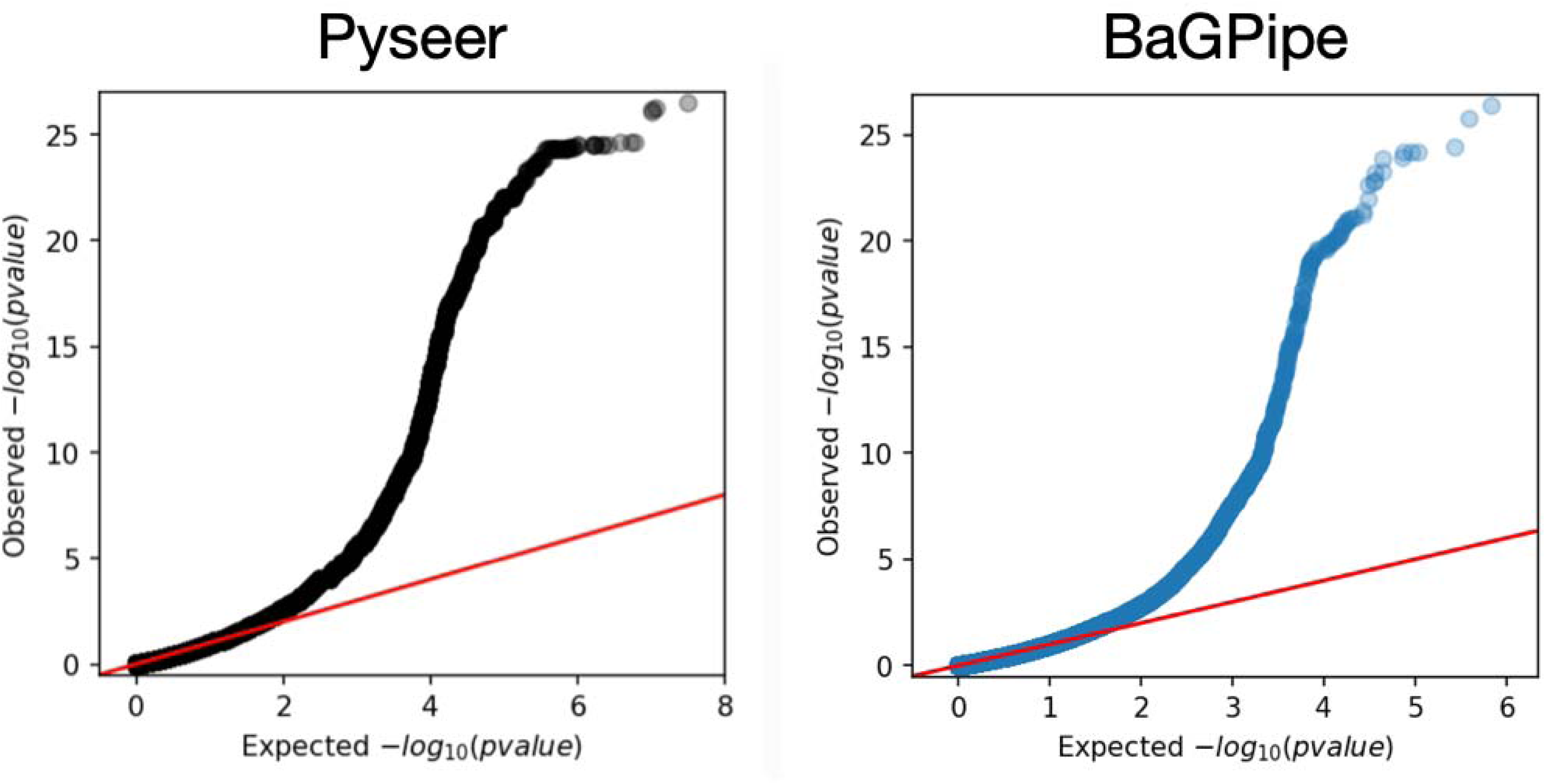
Comparison of the Q-Q plots from BaGPipe and from Pyseer on the Streptococcus pneumoniae dataset. The Q-Q plots show that observed −log_10_(p-values) are not inflated at low −log10(p-values) and there is an absence of any poorly controlled confounding population structure (they would appear as big “steps” deviating from the diagonal line). The position of the points being above the null hypothesis (the diagonal line) indicates significant k-mers/unitigs associated with penicillin resistance. The Q-Q plot produced from Pyseer was sourced from the Pyseer tutorial [29].

The subsequent significant unitig analysis conducted by BaGPipe identified 573 significant unitigs, in contrast to the 5,327 significant k-mers reported in the Pyseer tutorial. This notable reduction demonstrates the effectiveness of using unitigs to reduce sequence redundancy while retaining significant biological information. The significant unitigs were mapped to the reference genome, and a Manhattan plot was generated, showing two distinct peaks corresponding to the two penicillin-binding protein (pbp) genes: *pbp2x* and *pbp2b* (**Figure 4**). These are the hits found in the previous studies and they are known genes involved in penicillin resistance [29, 31, 32].

**Figure 4:**
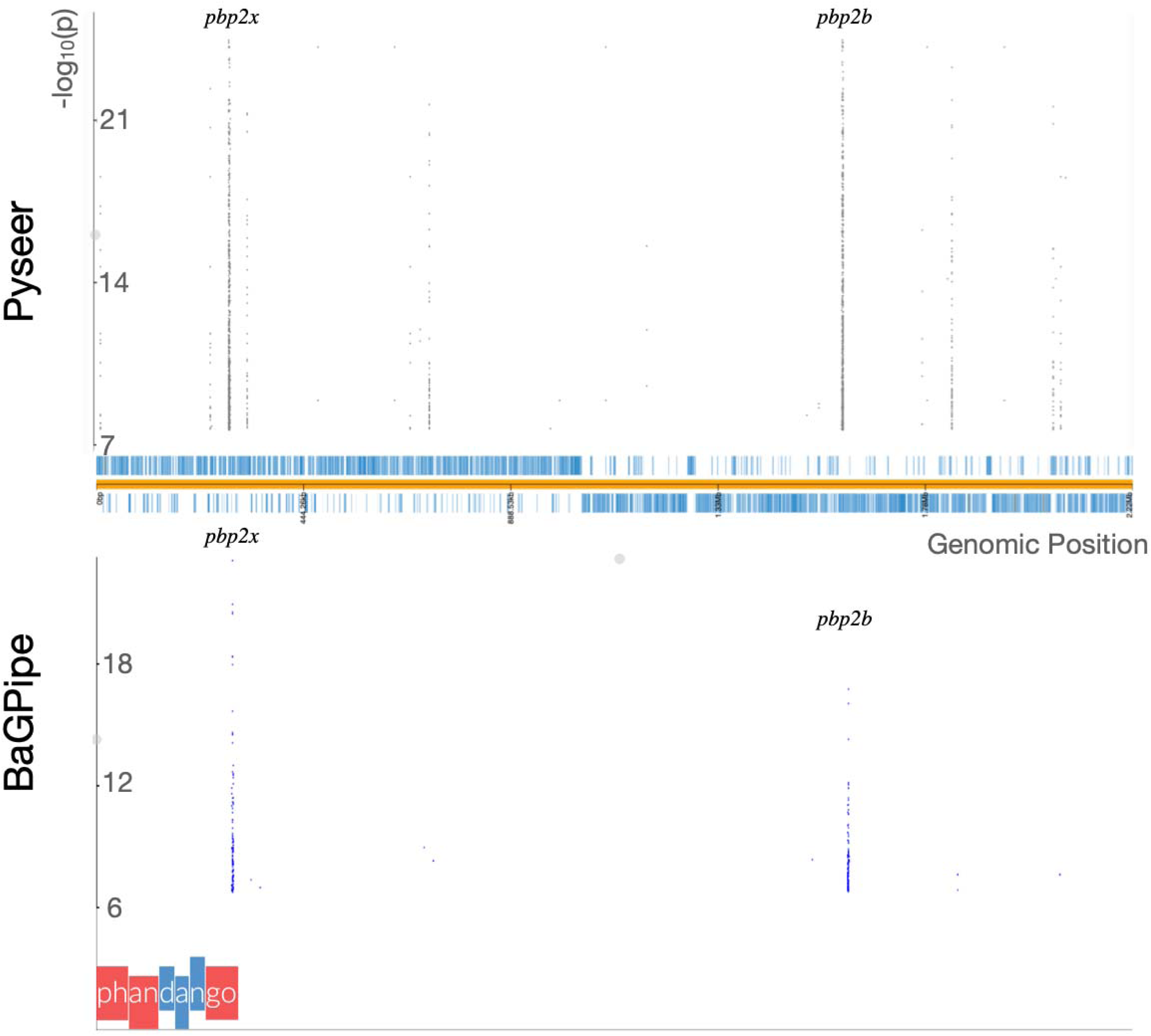
Comparison of the Manhattan plots from BaGPipe and from Pyseer on the Streptococcus pneumoniae dataset. The plot is again consistent with that in the Pyseer tutorial, showing two peaks, corresponding to the two pbp genes. The strongest peak represents the locus coding for the putative penicillin binding protein 2x (pbp2x); the second strongest peak represents pbp2b gene. The peaks present in the Pyseer plot but absent in the BaGPipe plot are likely to represent false positives associated with genes that have low minimum allele frequencies, as detailed in Figure 5. The Manhattan plot produced from Pyseer was sourced from the Pyseer tutorial [29] and the plots are visualised via Phandango [43].

Not all significant unitigs mapped to a single reference, underscoring the need for multiple high-quality reference genomes to ensure comprehensive annotation and reliable downstream analyses. The Manhattan plot generated by BaGPipe was consistent with that from the Pyseer tutorial, although certain peaks observed in the Pyseer tutorial’s results were absent in the BaGPipe’s results, indicating the presence of potential false positives in the former due to low minimum allele frequencies (MAF).

After annotating the significant unitigs, BaGPipe produced a list of "gene hits," which included unitigs that either resided within or were adjacent to specific coding regions. The gene-hit plot summarising these results closely matched that of the Pyseer tutorial (**Figure 5**). Key hits such as *penA* (*pbp2b*) and *pbpX* (*pbp2x*) were consistently identified across both analyses, demonstrating BaGPipe’s reliability. Interestingly, the *recR* gene, identified as significant in the Pyseer analysis, showed a lower significance in the BaGPipe-derived results. This discrepancy suggests that BaGPipe offers a more precise analysis, as *recR* may have been artificially associated in the original Pyseer analysis due to linkage with causal variants within overlapping regions of the *pbp* genes. *recR* encodes a protein involved in DNA repair, specifically the RecFOR pathway for homologous recombination [44], which may occasionally show spurious associations due to proximity to key resistance genes.

**Figure 5:**
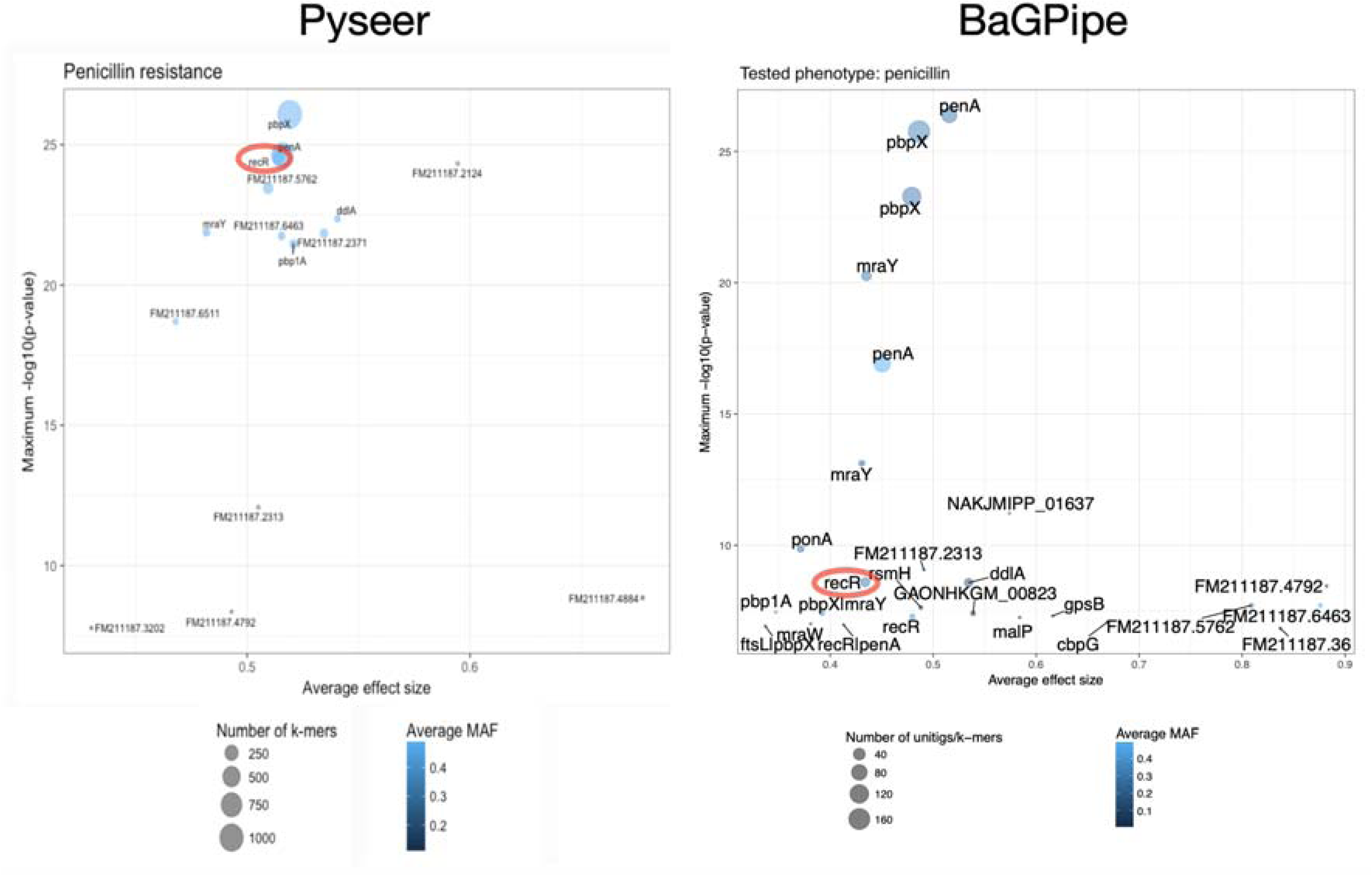
Comparison of the gene-hit plots from BaGPipe and from Pyseer on the S. pneumoniae dataset. The x-axis represents the average effect size, while the y-axis shows the maximum −log_10_(p-value), highlighting the statistical significance of gene associations. The size of the circular dots indicates the number of k-mers/unitigs involved in each association, providing insight into the genomic support for each hit. The colour scheme, ranging from lighter to darker shades of blue, illustrates the average MAF, with darker shades indicating a lower MAF. BaGPipe accurately replicated the findings from Pyseer, confirming penA (pbp2b) and pbpX (pbp2x) as the most significant gene hits. Red circles in the diagrams indicate recR, which Pyseer incorrectly identified as a major hit. In contrast, BaGPipe correctly depicted it as less significant. The gene-hit plot produced from Pyseer was sourced from the Pyseer tutorial [29].

Reproducing this analysis from raw reads, BaGPipe used approximately 191 hours of total CPU time, with a peak remote-access memory (RAM) usage of 3.5LGB, a mean RAM usage of 574□MB, and an overall run time of about 23 hours and 11 minutes. When excluding steps such as annotation, pangenome analysis, and phylogenetic analysis—assuming these are provided as input—the CPU time would be reduced to approximately 19 hours.

### Implementation of BaGPipe on a *S. aureus* Dataset

BaGPipe was further evaluated using high-quality sequences from 518 *Staphylococcus aureus* samples tested against seven antibiotics [33]. It successfully identified significant genetic associations with resistance phenotypes for multiple antibiotics. To cross-validate these findings, AMRFinderPlus [42] was employed to identify known antimicrobial resistance (AMR) genes. BaGPipe’s results were compared with AMRFinderPlus predictions, confirming the presence of established AMR genes, such as *gyrA*, *parC*, and *parE* for ciprofloxacin, and *ermA* for erythromycin (**Figure 6**). Additionally, BaGPipe identified gene hits absent in the AMRFinderPlus database, suggesting novel resistance mechanisms that warrant further investigation. Some genes identified by BaGPipe are known to confer resistance to other antibiotic classes, likely due to linkage disequilibrium within a shared mobile genetic element (MGE). This reflects the potential for cross-resistance or false-positive associations. However, these observations could also indicate mechanisms of resistance that are not yet fully understood, e.g. a regulatory gene. A detailed comparison of gene hits predicted by AMRFinderPlus and those identified by BaGPipe is summarised in **Table 1**. The counts of resistant and susceptible phenotypes for all eight antibiotics, along with Manhattan plots and gene hit plots produced from BaGPipe, are provided in the supplementary materials.

**Figure 6:**
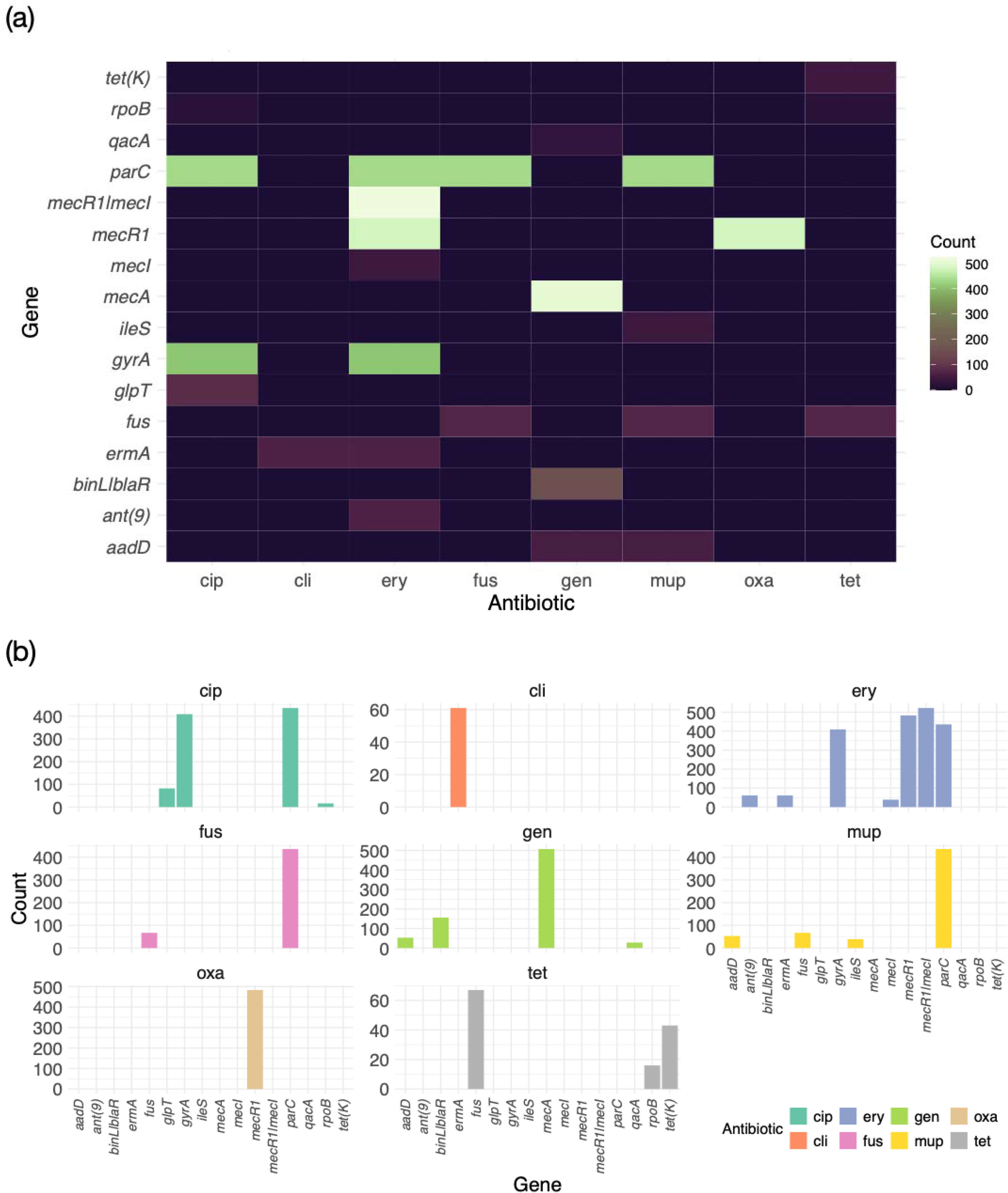
Frequency of matched gene hits from BaGPipe GWAS analysis and AMRFinderPlus prediction in the eight tested antibiotics for the S. aureus dataset. (a) Heatmap and (b) faceted bar plot of the frequency of matched genes across the eight tested antibiotics. The bars represent the counts of genomes in which gene hits identified by BaGPipe and predictions by AMRFinderPlus coincide, indicating the prevalence of known AMR genes within the tested strains. BaGPipe successfully identified known AMR genes in every tested antibiotic. cip, ciprofloxacin; cli, clindamycin; ery, erythromycin; fus, fusidic acid; gen, gentamicin; mup, mupirocin; oxa, oxacillin; tet, tetracycline.

**Table 1:**
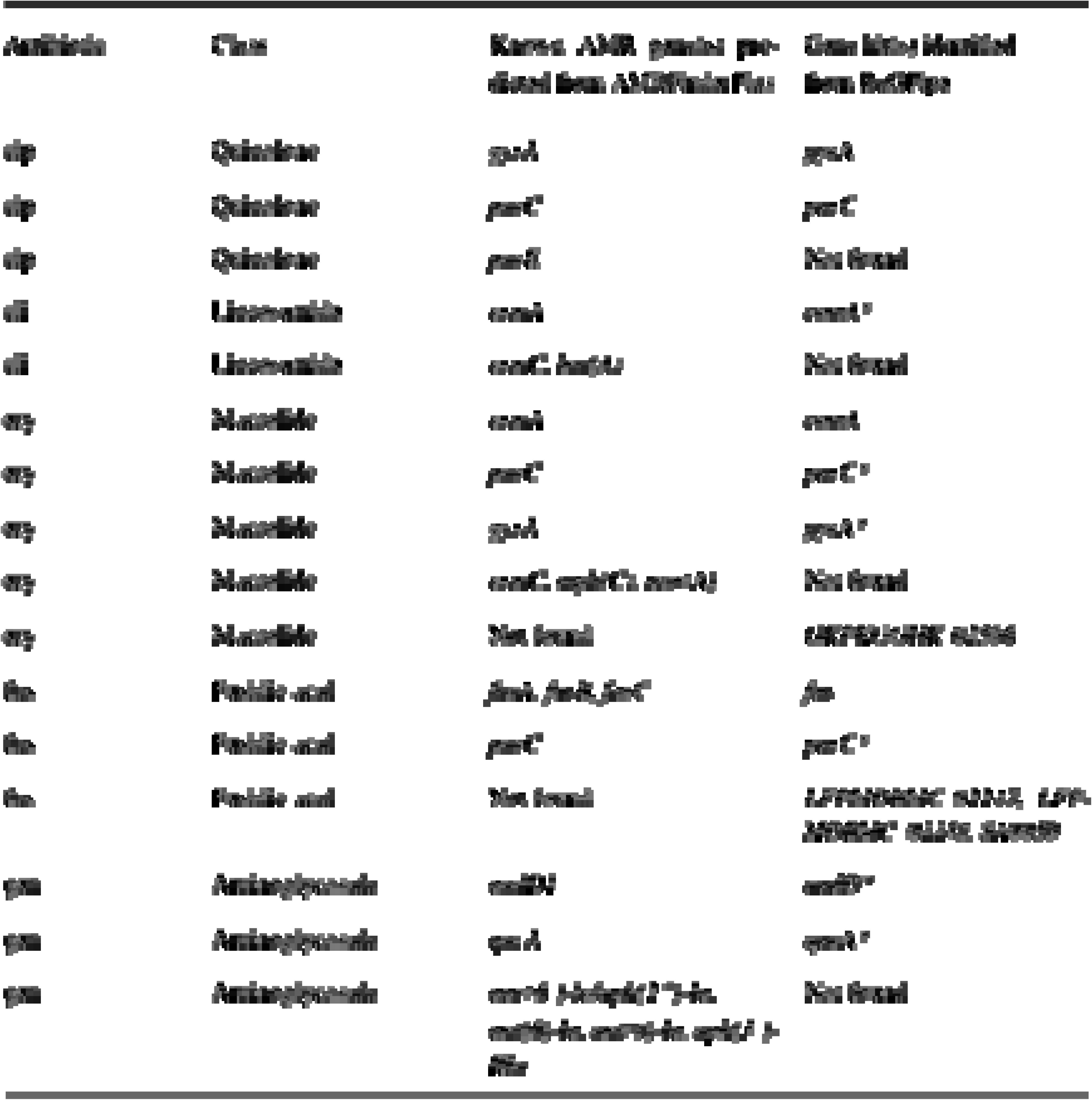
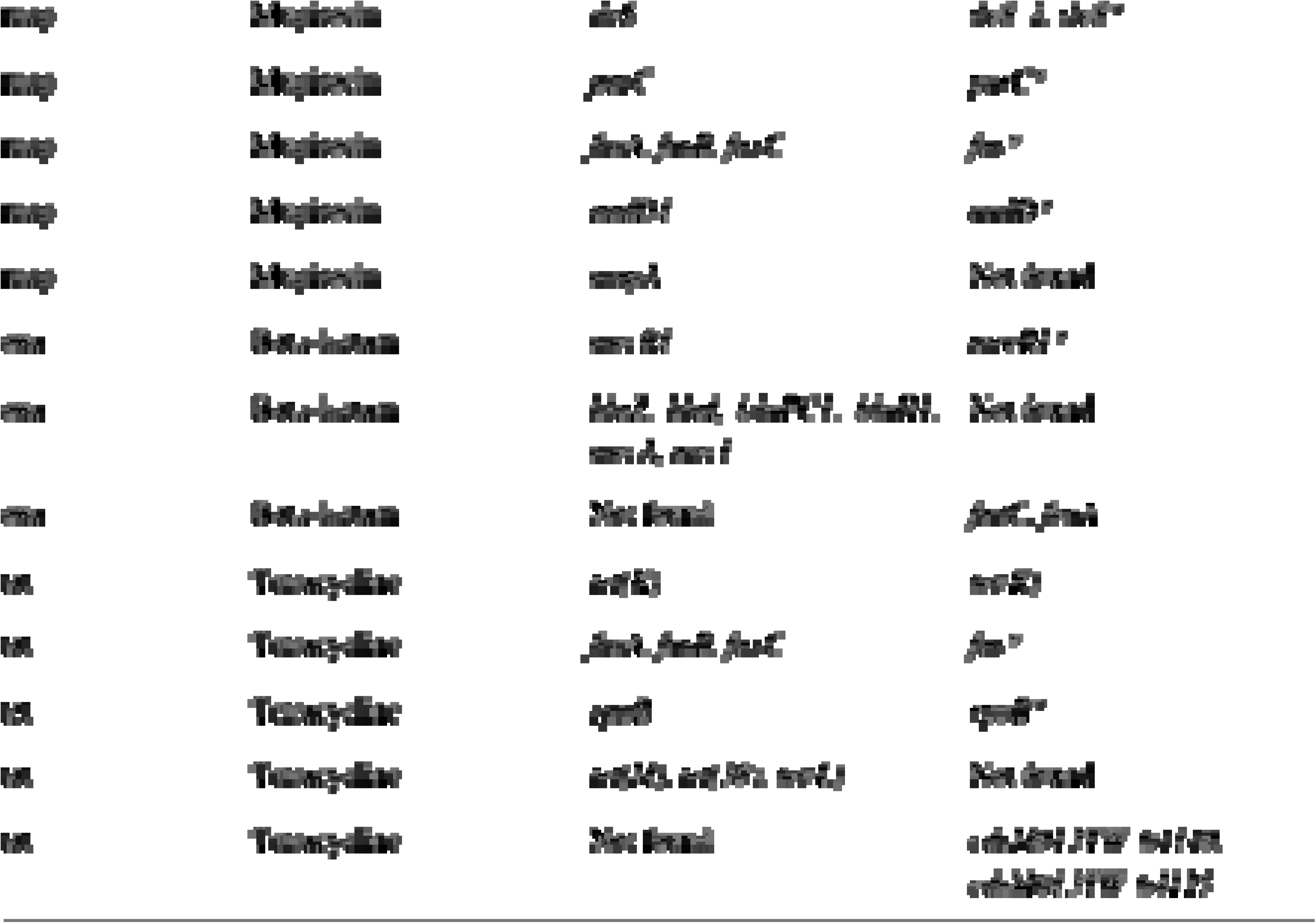
Gene hits predicted from AMRFinderPlus and identified from bacterial GWAS by BaGPipe for each of the eight tested antibiotics. For each antibiotic, the corresponding class was identified, and AMRFinderPlus was used to screen for known AMR genes within that class in its database. For example, for ciprofloxacin (cip), a quinolone, genes like gyrA, parC, and parE were predicted from the sequences. Conversely, BaGPipe’s targeted GWAS approach yielded hits such as gyrA and parC for ciprofloxacin, but not parE, suggesting that BaGPipe may be more sensitive and reflect the actual presence of AMR genes in the dataset more accurately. There are some gene hits identified by BaGPipe that are not predicted from the known database; these hits are manifested as identifiers extracted from GFF files during annotation. *: Not significant gene hit. cip, ciprofloxacin; cli, clindamycin; ery, erythromycin; fus, fusidic acid; gen, gentamicin; mup, mupirocin; oxa, oxacillin; tet, tetracycline.

## Discussion

BaGPipe addresses the challenges associated with analysing large-scale bacterial datasets by integrating a cutting-edge workflow management tool, Nextflow [30], along with a modular design that allows researchers to efficiently conduct end-to-end analyses. This pipeline offers an accessible solution to a major bottleneck in bacterial genomics: performing reproducible and efficient GWAS on large datasets with diverse computational requirements.

BaGPipe successfully reproduced a published *S. pneumoniae* study [29, 31, 32], validating its reliability for bacterial GWAS. The analysis was reproduced efficiently, demonstrating optimised computational resource consumption. Its successful application to a *S. aureus* dataset further demonstrated versatility across different microbial contexts. However, the dataset composition can influence association signals, as seen in the *S. aureus* analysis, where certain antimicrobial resistance (AMR) genes (*parC* for fusidic acid and mupirocin, *mecA* for gentamicin) were identified despite not being direct resistance determinants. This likely reflects inherent biases in the dataset, including linkage disequilibrium, clonal population structure, resistance genes co-localised on the same MGE, and complex regulatory interactions that modulate gene expression. Additionally, the proportion of resistant versus susceptible isolates can affect gene associations. Nevertheless, cross-validation with AMRFinderPlus [42] confirmed the presence of established AMR genes and identified additional gene hits absent in existing AMR databases. While these loci are not confirmed resistance genes, they represent promising leads for further experimental investigation, as these associations do not necessarily indicate false positives. For instance, mutations in *rpoB*, initially linked to rifaximin resistance, were recently shown to confer daptomycin resistance in *Enterococcus* faecium [19]. These findings highlight both BaGPipe’s sensitivity and robustness in hypothesis generation for discovering unexplored genetic mechanisms.

BaGPipe’s advantage is rooted in its flexibility, versatility and adaptability. Its modular design not only integrates seamlessly across diverse research scenarios but also allows users to incorporate their preferred methods for annotation, tree construction and other analyses. By automating resource management and dynamically allocating computational resources, BaGPipe simplifies large-scale analyses without the need for manual adjustments.

In terms of computational resource usage, we believe BaGPipe is designed to be user-friendly and efficient, handling moderately sized datasets with ease, as demonstrated by its low CPU time usage, modest remote-access memory (RAM) requirements, and overall runtime of 23 hours in the *S. pneumoniae* study. It is worth noting that these metrics reflect the datasets used in this study and should be viewed as indicative, as computational performance may vary depending on the input size, complexity, and the specific steps included in each given analysis.

The benchmarking of BaGPipe against other existing tools is another area that remains limited. Due to the scarcity of comprehensive bacterial GWAS pipelines, an exhaustive comparative analysis was challenging. As the field evolves, further benchmarking efforts that include newer tools will be necessary to fully evaluate and demonstrate BaGPipe’s competitive edge.

Future directions for BaGPipe involve the expansion of its capabilities to include additional genotype formats, such as single nucleotide polymorphisms (SNPs) and gene presence/absence, as well as the incorporation of continuous variables for phenotypes. With continued improvements and community-driven development, BaGPipe has the potential to become an accessible and indispensable tool for researchers in microbial genomics, fostering new discoveries and deepening our understanding of bacterial genetics.

In conclusion, BaGPipe is a powerful, flexible, and efficient pipeline for conducting bacterial GWAS with the well-established Pyseer framework. We show it is capable of reliably handling complex datasets, reducing redundancy, and replicating or uncovering both known and novel genetic associations related to antibiotic resistance. BaGPipe lowers the barriers to performing bacterial GWAS and offers a comprehensive, reproducible framework, vital to better understand the genetic basis of AMR and bacterial traits.

## Supporting information

Supplementary Main File

## Abbreviations

AMR: Antimicrobial Resistance
DRM: Distributed Resource Manager
FaST-LMM: Factored Spectrally Transformed Linear Mixed Model
GWAS: Genome-wide Association Study
HPC: High-performance Computing
Indel: Insertions and Deletions
LSF: Load Sharing Facility
MAF: Minimum Allele Frequency
MDS: Multidimensional Scaling
MGE: Mobile Genetic Element
MRSA: Methicillin-Resistant *Staphylococcus aureus*
PCA: Principal Component Analysis
Q-Q plot: Quantile-Quantile plot
RAM: Remote-Access Memory
SNP: Single Nucleotide Polymorphism
VCF: Variant Calling Format

## Author statements

### Author contributions

D.A., W.R.S., E.M.H. designed the study; K.W. built the pipeline with input from W.R.S. and undertook the bioinformatic analyses with contribution from D.A. and W.R.S.; W.R.S., D.A., and E.M.H. supervised the study; B.B. curated and provided the *S. aureus* dataset; S.R., M.S.T., E.T., S.J.P. contributed significantly to data generation and analysis of the original study which produced the *S. aureus* dataset; K.W. wrote the first draft of the manuscript; D.A., W.R.S., E.M.H., K.W. edited the manuscript. All authors had access to the data and read, contributed and approved the final manuscript.

## Acknowledgements

We would like to appreciate John Lees and his team for making Pyseer and its tutorial available. We would also like to appreciate Francesc Coll for their encouragement and feedback.

