## Supplementary Main File for "BaGPipe: an automated, reproducible, and flexible pipeline for bacterial genome-wide association studies"

**Supplementary Table 1: Current prominent methods in overcoming bacterial GWAS challenges.** Most tools demand additional software and expertise from the user to execute complex analyses. BH adjustments, Benjamin-Hochberg adjustments; FaST-LMM, Factored Spectrally Transformed Linear Mixed Models; MDS, Multidimensional Scaling; PCA, Principal Component Analysis.

| Software | Type of Approach | Population Structure Adjustment | Recombination Rate Adjustment | Multiple Testing Adjustment | Polygenicity |
|---|---|---|---|---|---|
| Scoary | Phylogenetic | Pairwise comparisons | No | Bonferroni and BH adjustments | Partially |
| TreeWAS | Phylogenetic | Phylogenetic inference | Yes | Bonferroni and False Discovery Rate adjustments | Yes |
| Bugwas | Non-phylogenetic | PCA | No | Bonferroni correction | Yes |
| HAWK | Non-phylogenetgic | PCA | No | Bonferroni and BH adjustments | Partially |
| SEER | Non-phylogenetic | Distance matrix derived from k-mers with MDS application | Yes | Bonferroni correction and Permutation testing | Yes |
| Pyseer | Non-phylogenetic | Fixed effects via MDS in phylogenetic regression and FaST-LMM with random effects | Yes | Threshold determined through hashing k-mers | Yes |
| Kover | Machine learning | No | No | Not affected | No |
| Phenotype-Seeker | Machine learning | Distance matrix | Not specified | Not specified | Not specified |

**Supplementary Table 2: Metadata of all 520 *Staphylococcus aureus* assemblies including their ERS accession numbers.** Attached separately as an Excel file.

**Supplementary Table 3: Data for Top Five Species Plot for MultiQC report produced from BacQC (Kraken2 and Bracken) on the *Staphylococcus aureus* assemblies.** Attached separately as an Excel file.

**Supplementary Table 4: Number of resistant and susceptible *Staphylococcus aureus* isolates for each tested antibiotics.** Attached separately as an Excel file.

**Supplementary 5: Reference genome assemblies used in the implementation of BaGPipe in the analysis with the *Staphylococcus aureus* dataset.** Assembly accession number are shown here and these can be searched on the NCBI Assembly database.

Reference genome assemblies used by BaGPipe in its application on the *S. aureus* dataset:

1. GCA_000009645.1_ASM964v1
2. GCA_000237125.3_ASM23712v3
3. GCA_000953255.1_Staphylococcus_aureus_Sa_ILRI_217
4. GCA_000011265.1_ASM1126v1
5. GCA_000017085.1_ASM1708v1
6. GCA_001611405.1_ASM161140v1
7. GCA_000009585.1_ASM958v1
8. GCA_000412775.1_ASM41277v1

**Supplementary 6: Top five species plot for the quality control of the *Staphylococcus aureus* genomes.** The plot was produced by the BacQC pipeline (Kraken2 and Bracken) and it clearly showed two outliers that had been species contamination. The two isolates were removed from the analyses. The raw data that produced this plot can be found as a supplementary Excel table (**Supplementary Table 3**).

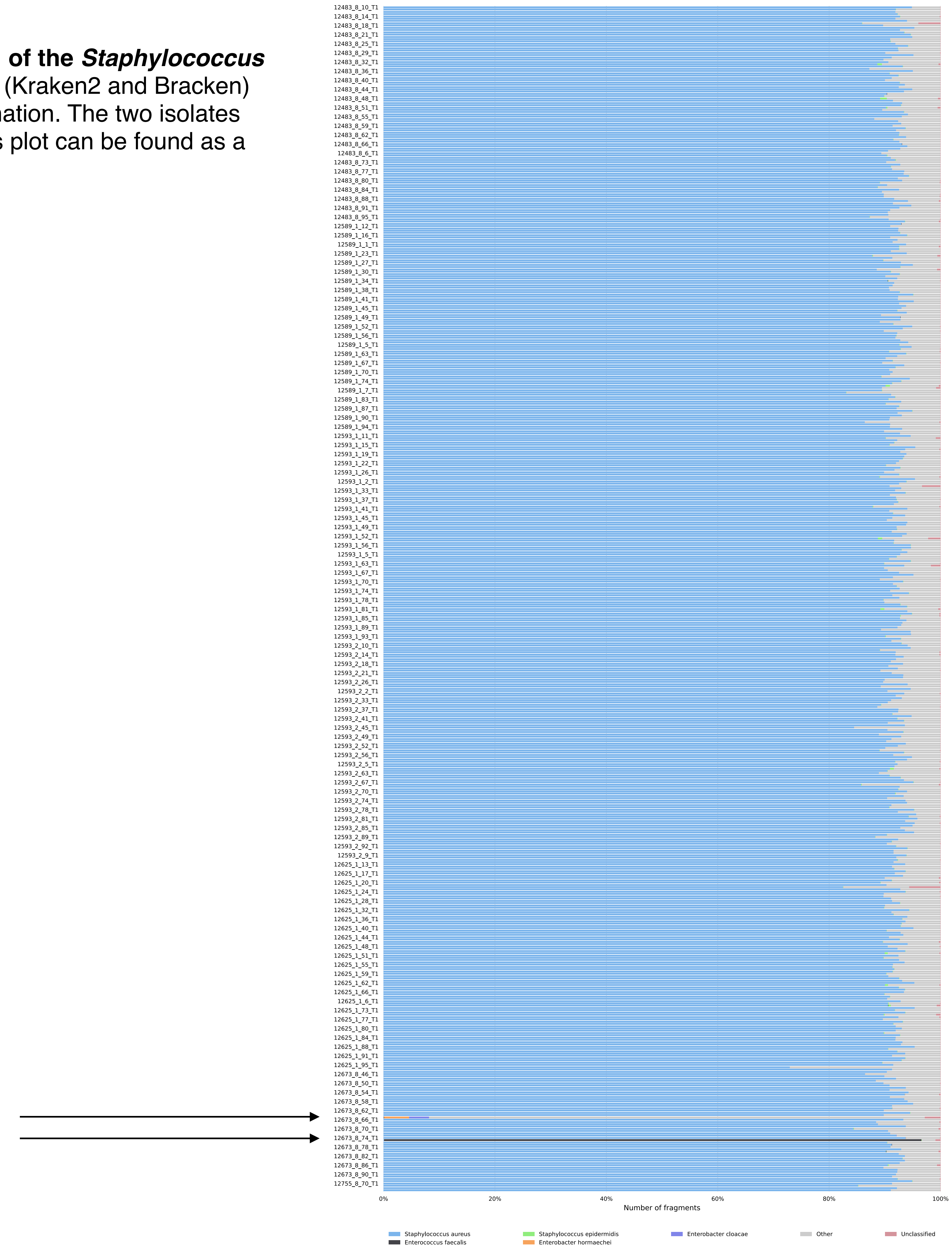

**Supplementary 7: Manhattan plot raw files produced by BaGPipe for all 8 GWAS analyses in the *Staphylococcus aureus* dataset.** These can be viewed using Phandango, with the addition of a reference GFF file. These can be found on the GitHub repository of BaGPipe (https://github.com/sanger-pathogens/BaGPipe). Below shows one example of such Manhattan plot visualisation using Phandango.

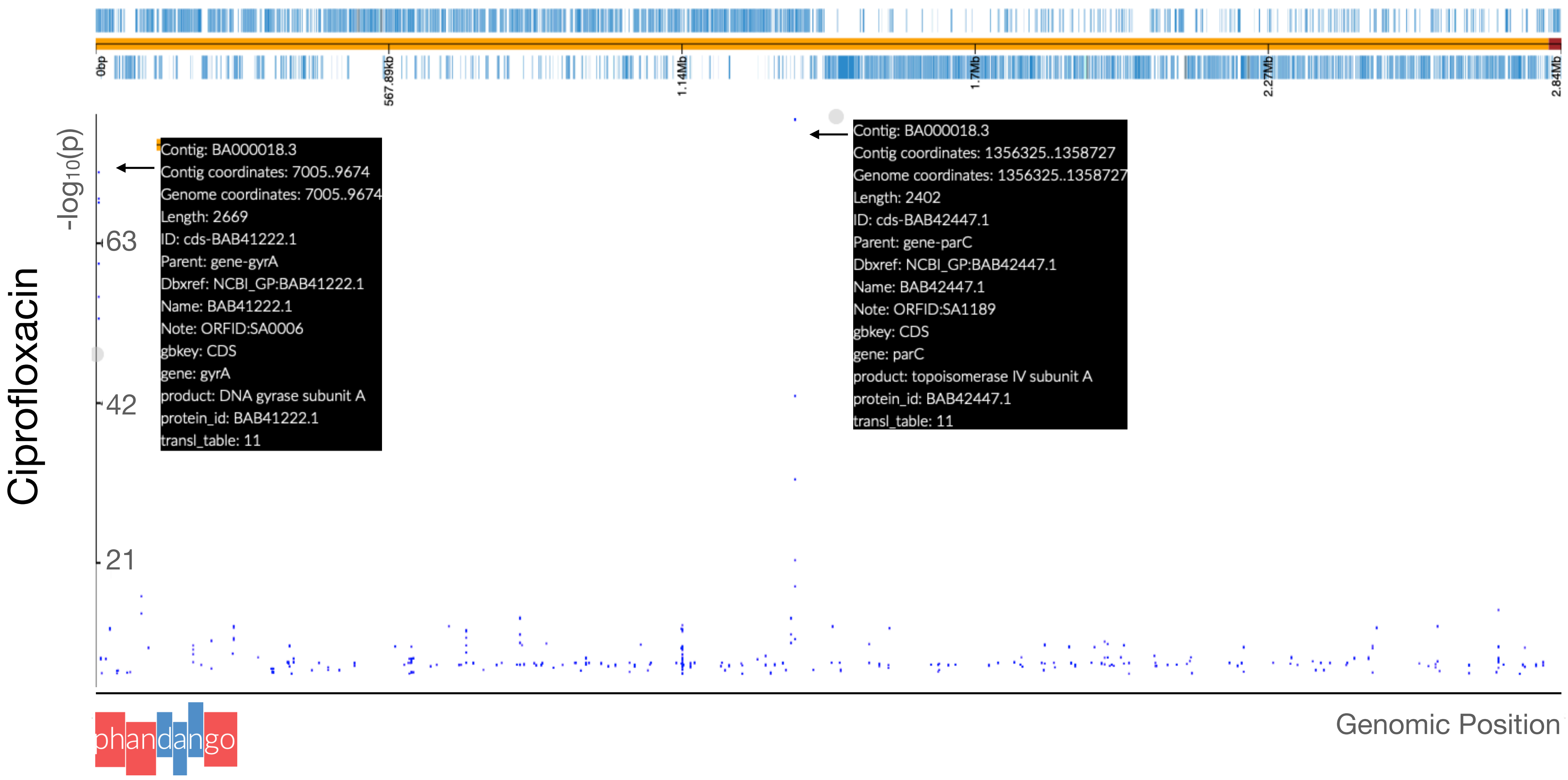

**Supplementary 8: Annotated unitigs and Gene Hit Plots produced by BaGPipe for all 8 GWAS analyses in the *Staphylococcus aureus* dataset.** These can be found on the GitHub repository of BaGPipe (https://github.com/sanger-pathogens/BaGPipe).
